# BAP31 regulated polarization of macrophages through C/EBP β in cutaneous wound healing

**DOI:** 10.1101/2020.05.14.095554

**Authors:** Qing Yuan, Bo Zhao, Yu-hua Cao, Jia-cheng Yan, Li-jun Sun, Xia Liu, Yang Xu, Xiao-yu Wang, Bing Wang

**Affiliations:** College of Life Science and Health, Northeastern University, #195 Chuangxin Road, Hunnan Xinqu. Shenyang, Liaoning 110169, China

**Keywords:** BAP31, BMDM, C/EBP β, Cutaneous wound healing, M2 polarization

## Abstract

The functions carried out by macrophages are essential in the processes of repairing skin injury. However, the mechanism of the M2 macrophage and its role in cutaneous wound healing remain elusive. B cell receptor associated protein 31 (BAP31) plays an important role in the immune system, and its function in connection with macrophages has yet to be determined. The present study demonstrates that the process of cutaneous wound healing slowed down in bone marrow-specific BAP31 knock down Lyz2-cre-BAP31^flox^/^flox^ mice. In addition, further studies show that various kinds of macrophage M2 polarization related factors were regulated by BAP31. Among these molecules C/EBP β was significantly affected. However, IL-4 but not IFN-γ, is able to recover the expression levels of C/EBP β and its downstream transcript factors induced by BAP31. Then, we demonstrated that BAP31 regulated macrophage M2 polarization by negative regulation of IL-4Rα and positive influence on Egr-2 to affect C/EBP β. Our findings reveal a novel mechanism of BAP31 in regulating M2 macrophage, and provide novel targets for the prevention and treatment of chronic wounds.

## Introduction

Cutaneous wound healing is a physiological response of skin to external injury, consists of four stages namely hemostasis, inflammation, epithelial remodeling and remodeling. The function of macrophages is closely related to the repair of skin injury (Esposito *et al*, 2019; Kim *et al*, 2019). Previous studies have shown that macrophages are important in regulating of host defense in organisms (Li *et al*, 2019). In order to meet the different needs of the body, macrophages can be transformed into different functional phenotypes under different stimulating factors, a process called polarization. Resting macrophages (M0) can polarize to form different phenotypes, proinflammatory phenotype (M1) and anti-inflammatory phenotype (M2) under different influencing factors (Li *et al*, 2018; Miao *et al*, 2017). M2 macrophages participate in the whole process of repairing skin injury (Du Y *et al*, 2018). The balance of cytokines and inflammatory factors in tissue repair is regulated by M2 macrophages (Zhang *et al*, 2018).

B-cell receptor-related protein 31 (BAP31) is an indispensable transmembrane protein in the endoplasmic reticulum (ER) with a molecular weight of 28 kDa (Wang *et al*, 2008). BAP31 is evolutionarily conservative and widely expressed in various tissues that mainly involved in the proliferation, invasion and metastasis of cancer cells, as well as in the process of apoptosis (Machihara & Namba, 2019). Previous studies in our laboratory have shown that BAP31 can affected the function of fat and the occurrence of postsynaptic diseases (Jia CC *et al*, 2018; Xu JL *et al*, 2018). Moreover, involving immunity, BAP31 can regulate T cell function through TCR signaling pathway, and influencing the function of macrophages (Fortier A *et al*, 2007; Niu *et al*, 2017). BAP31 deficiency upregulates LPS-induced proinflammatory cytokines in BV2 cells and mice by upregulating the protein level of IRAK1 in neuroinflammation (Liu *et al*, 2019). Therefore, we consider that BAP31 may affect the polarization of macrophages and the cutaneous wound healing.

There are several key transcription factors clearly associated with macrophage polarization as STAT family, peroxisome proliferator-activated receptor (PPAR), cAMP response element binding protein (CREB)-CCAAT/enhancer binding protein (C/EBP), hypoxia-inducible factors (HIF), NF-*κ*B and IFN regulatory factors (IRF). Several genes associated with M2 macrophage phenotype (Arg1, CD206 and Ym1) are regulated by STAT-6 activity in the presence of IL-4/IL-13, another factor required for M2 polarization is PPAR-c, which is also inducible by IL-4/IL-13. Additionally, C/EBP β regulates many M2-related genes and CREB-C/EBP β activity is required for wound healing which is an M2 macrophage function (Funes *et al*, 2018; Saradna *et al*, 2018). CCAAT enhancer binding proteins β (C/EBP β) is a transcription factors, the important member of the C/EBPs family. C/EBP β is involved in the regulation of cell proliferation and differentiation, tumorigenesis and apoptosis, and hematopoietic cell formation (Ramji DP & Foka P, 2002; Shackleford *et al*, 2011; Zahnow, 2009). Nowadays it is acknowledged that C/EBP β is important in M1 and M2 macrophage polarization, high expression of C/EBP β in M1 macrophages, and through changing TLR4-HMGB1/C/EBP β pathway promotes the M2 polarization (Veremeyko *et al*, 2018). The expression of C/EBP β is influenced by early growth response gene-2 (Egr-2), a member of a family of zinc finger proteins, has been found to play a critical role in hindbrain development and myelination in the peripheral nervous system, and also regulates the development of B cells, T cells, and macrophages (Li *et al*, 2011; Zhu *et al*, 2008).

This study reports that BAP31 conditional knockout can slow down the process of cutaneous wound healing, and inhibited M2 macrophage polarization via the transcription factors C/EBP β. Moreover, BAP31-deficiency reduced the expression of Egr-2 which is the regulatory factors for C/EBP β, but enhanced the expression of IL-4Rα that caused the recovery of the expression of C/EBP β downregulated by BAP31-deficiency in IL-4-stimulated macrophages. These novel roles of BAP31 in macrophages will provide a potential clue for the treatment of chronic wounds.

## Results

### BAP31 knockout retarded the cutaneous wound healing in mice

In order to investigate the function of BAP31 in immune system we generated BAP31 conditional knockout mice (Niu *et al.*, 2017). The BAP31 floxed mice (BAP31^flox/flox^) were bred with Lyz2-Cre transgenic mice to induce specific deletion of this gene in bone marrow macrophages. Bone marrow cells of Lyz2-cre BAP31^flox/flox^ (BAP31^-/-^) and BAP31^flox/flox^ (BAP31^+/+^) mice were separated and induced with the supernatant of L929 cells to let them differentiate into Bone Marrow Derived Macrophages (BMDMs). FACS analysis revealed that the proportion of F4/80 and CD11b double positive cells from both mice were over 98% (**Fig 1A**). The expression of BAP31 in BMDMs was analyzed by RT-qPCR and Western analysis. The results showed that BAP31 was knocked down about 80% both in protein and mRNA of BMDMs (**Fig 1B and C**). Therefore, Lyz2-cre BAP31^flox/flox^ mice can specifically knockdown the expression of BAP31 in BMDMs. To assess the functions of BAP31 in immune related wound repair, we created a murine model of cutaneous wound healing in BAP31^-/-^ and BAP31^+/+^ mice. The wound area was measured every 3 days, and found that the wound contraction and healing processes of BAP31^-/-^ mice were slowed down (**Fig 1D and E**). Morphological analysis via hematoxylin and eosin (H&E) staining of the wound tissue sections at 14 days after injury revealed that the skin tissue of BAP31^-/-^ mice were not completely repaired compared with the control mice (**Fig 1F**). Ym1 was an essential anti-inflammatory gene of wound healing (Finley PJ *et al*, 2016). Immunofluorescence staining revealed that Ym1 in cutaneous wound tissue of BAP31^-/-^ mice was significantly reduced compared with that in the BAP31^+/+^ mice (**Fig 1G**). These results implied that specific deletion of BAP31 could slowed down the process of cutaneous wound healing in mice.

**Figure 1.**
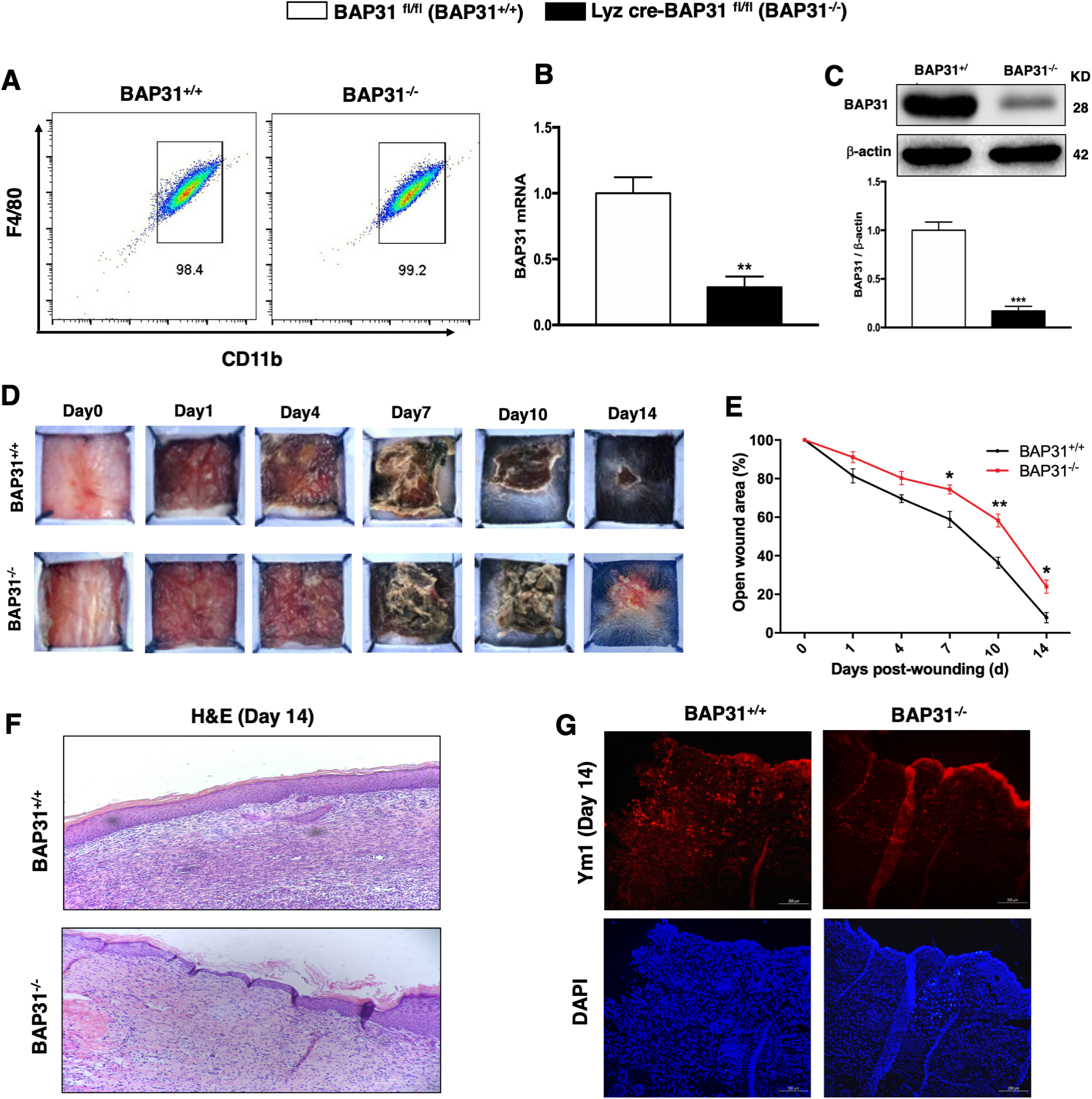
BAP31 knockout retarded the cutaneous wound healing in mice. A FACS analysis BMDMs which were extracted from BAP31^-/-^ and BAP31^+/+^ mice using FITC-CD11b and PE-F4/80. B RT-qPCR analysis of BAP31 mRNA from BMDMs of BAP31^-/-^ and BAP31^+/+^ mice. C Western-blot analysis of BAP31 protein expression in BMDMs from BAP31^-/-^ and BAP31^+/+^ mice, histograms showed BAP31 relative change. D The skin wounds were measured in 0, 1, 4, 7, 10, 14 days after injure by camera. E The injury areas were measured in 0, 1, 4, 7, 10, 14 days after injure by Image J software. F The skin tissue images of H&E staining at day 14 after injure. G Images of fluorescent staining of skin tissue at day 14 after injure with Ym1 and DAPI. Data information: In (B, C, E) data presented as mean ± SD. n>3. *P < 0.05 versus BAP31-/^-^, **P < 0.01 versus BAP31^-/-^, ***P < 0.001 versus BAP31^-/-^.

### BAP31 affected the expression of M2 macrophage related transcription factor C/EBP β

The above information prompted us to examine whether BAP31 is involved in the regulation of macrophage M2 polarization. First, we investigated M2 polarization related transcription factor families, IRFs, C/EBPs and PPAR-γ, by RT-qPCR analysis in BMDMs (Wang *et al*, 2014). It is noteworthy that the expression of IRF1, IRF2, IRF5, and C/EBP β were decreased after BAP31 knockout, and C/EBP β was affected significantly (**Fig 2A and B**). In addition, FACS analysis revealed that the expression of C/EBP β was significantly decreased in BMDMs of BAP31^-/-^ mice (**Fig 2C**). Similarly, the protein expression levels of C/EBP β was also down-regulated in BAP31^-/-^ mice (**Fig 2D**). Conversely, after BAP31 overexpression in RAW264.7 cells, the protein and mRNA expression of C/EBP β were increased (**Fig EV2A and B**). These results suggested that BAP31 regulated the expression of C/EBP β in macrophages. Next, we studied the expression of M2 macrophage markers regulated by these transcription factors, consisted of Arg-1, Fizz-1, Ym1 and CD206 of BMDMs in BAP31^-/-^ and BAP31^+/+^mice. RT-qPCR analysis revealed that the expression of Arg-1, Fizz-1, Ym1 and CD206 were significantly decreased after the knockout of BAP31 (**Fig 2E**). FACS analysis revealed a profound reduction of two major M2 macrophage related molecules, CD206 and Ym1 in the BMDMs of BAP31^-/-^ (**Fig 2F**). Western-blot analysis showed that the expression levels of Arg-1 and Ym1 dramatically decreased after BAP31 knockout (**Fig 2G**). In addition, from the BAP31 overexpressed RAW264.7 cells (**Fig EV1A and B**), we found the mRNA expression levels of M2 signature genes Arg-1, Fizz-1, Ym1 and CD206 increased (**Fig EV1C**), indicating that BAP31 is involved in M2 polarization of macrophages. Taken together, these findings indicated that BAP31 regulated M2 macrophage and its related transcription factors such as C/EBP β.

**Figure 2.**
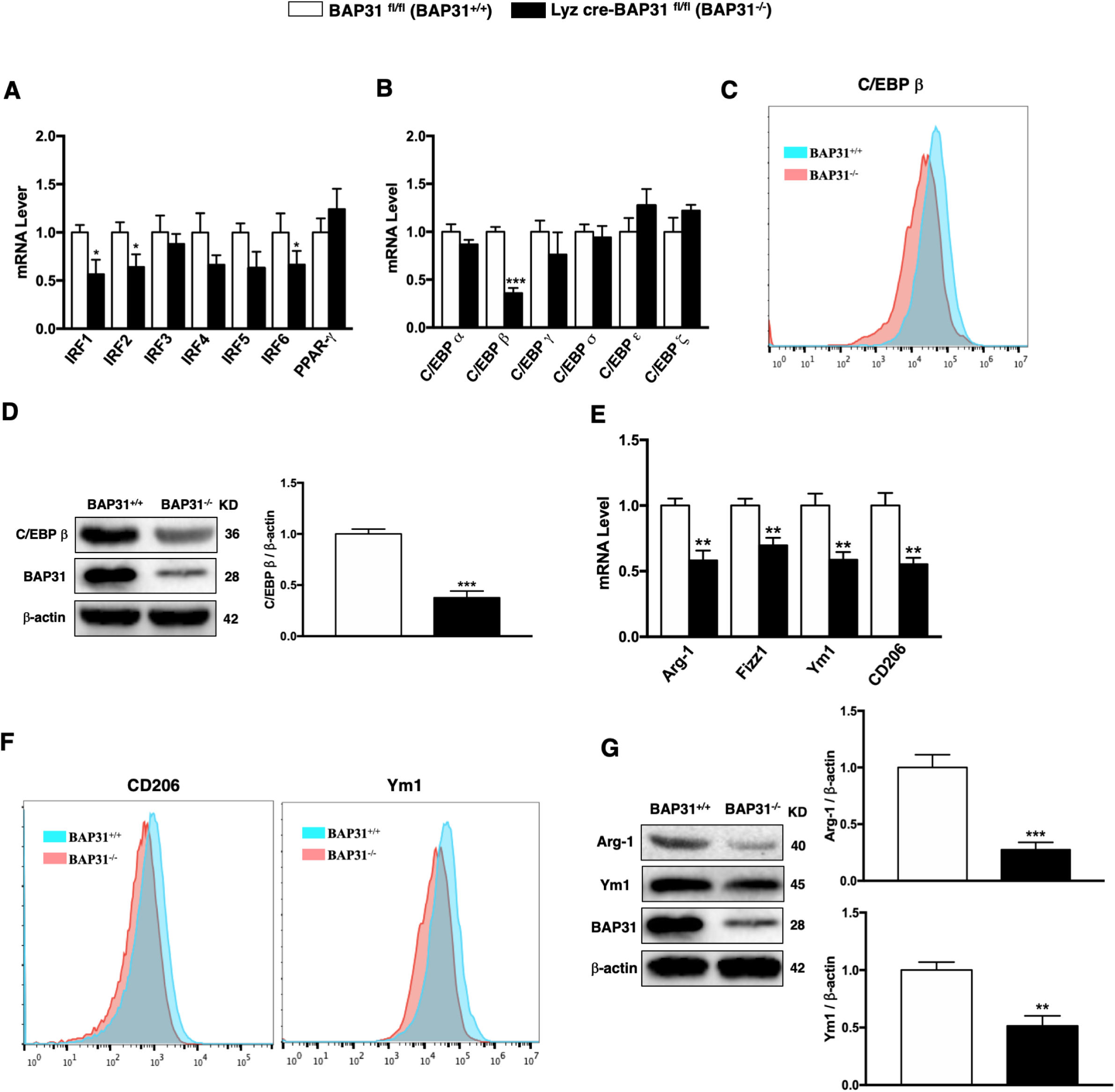
BAP31 affected the expression of M2 macrophage related transcription factor C/EBP β. A, B RT-qPCR analysis IRFs, C/EBPs, PPARγ mRNA from BMDMs of BAP31^-/-^ and BAP31^+/+^ mice. C FACS analysis the expressions of C/EBP β in BMDMs of BAP31^-/-^ and BAP31^+/+^ mice. D Western-blot analysis of C/EBP β protein expressions in BMDMs of BAP31^-/-^ and BAP31^+/+^ mice, histograms showed the relative changes. E RT-qPCR analysis Arg-1, Fizz1, Ym1, CD206 mRNA from BMDMs of BAP31^-/-^ and BAP31^+/+^ mice. F FACS analysis the expressions of Ym1and CD206 in BMDMs of BAP31^-/-^ and BAP31^+/+^ mice. G Western-blot analysis of Arg-1 and Ym1 in BMDMs of BAP31^-/-^ and BAP31^+/+^ mice, histograms showed the relative changes. Data information: In (A, B, D, E, G) data presented as mean ± SD. n>3. *P < 0.05 versus BAP31^-/-^, **P < 0.01 versus BAP31^-/-^, ***P < 0.001 versus BAP31^-/-^.

### IL-4 recovered the down regulation of C/EBP β induced by BAP31 in BMDMs

Literitures reported that C/EBP β played an important role both in the M1 macrophages and M2 macrophages (Lamkin *et al*, 2016; Ruffell D *et al*, 2009), therefore we added corresponding stimulation to induce M0 macrophages polarize to M1 and M2 macrophages, to further elucidate the impact of BAP31 on C/EBP β (Sahu *et al*, 2017). After seven days of culture, the BMDMs from BAP31^-/-^ and BAP31^+/+^ mice were stimulated by IFN-γ (20 ng/mL) and IL-4 (20 ng/mL) for 24 h respectively. Then, proteins of the above samples were collected for Western-blot analysis. We found that the expression of C/EBP β and its downstream factors Cox-2, Arg-1 were down-regulated in BAP31^-/-^ group. After stimulated by IFN-γ, the down regulation induced by the deletion of BAP31 keeps unchanged. Unexpectedly, after stimulated by IL-4, the down regulation of C/EBP β, Cox-2 and Arg-1 induced by BAP31 recovered to the original levels (**Fig 3A**). The results of RT-qPCR analysis also proved that the expression of C/EBP β, Cox-2 and Arg-1 significantly decreased after BAP31 knockdown in unstimulated and stimulated by IFN-γ in BMDMs, but after stimulated by IL-4, the expression of them were remained unchanged (**Fig 3B-F**). Similarly, the FACS results revealed that, the C/EBP β was down-regulated in BAP31^-/-^ BMDMs, but returned back after stimulated by IL-4 (**Fig 3G**). These results suggested that IL-4 stimulation can diminish the effects on the expression of C/EBP β induced by BAP31 in BMDMs.

**Figure 3.**
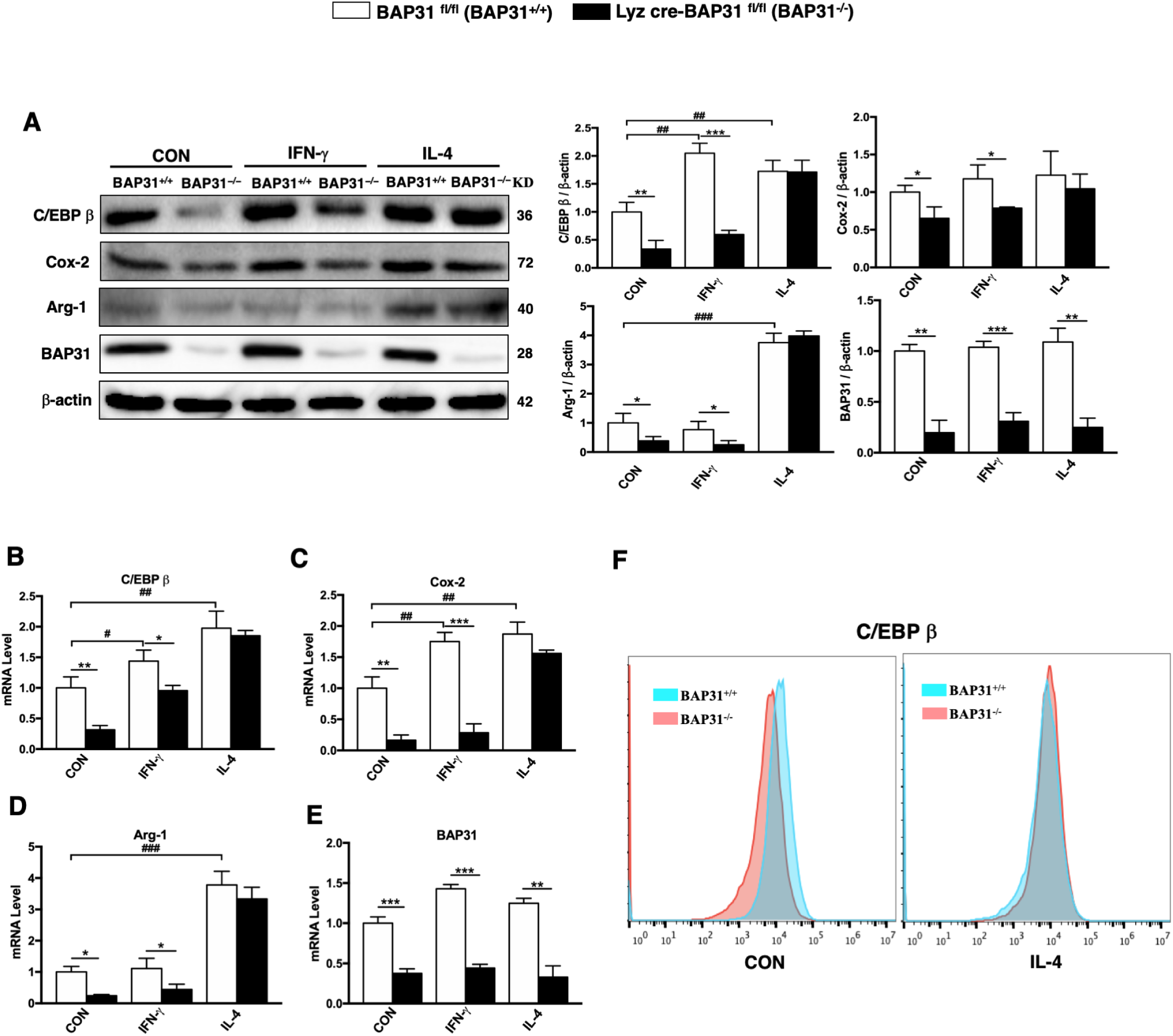
IL-4 recovered the down regulation of C/EBP β induced by BAP31 in BMDMs. A Western-blot analysis of C/EBP β, Cox-2 and Arg-1 protein expressions in BMDMs of BAP31^-/-^ and BAP31^+/+^ mice, samples were divided into con group, IFN-γ (20 ng/ml) stimulated group and IL-4 (20 ng/ml) stimulated group, both stimulated for 24 h. The histograms showed the relative changes. B-E RT-qPCR analysis the C/EBP β, Cox-2 and Arg-1 mRNA of the above cells. (**F**) FACS analysis the expressions of C/EBP β in BMDMs of BAP31^-/-^ and BAP31^+/+^ mice in IL-4 stimulated group and con group. Data information: In (A-E) data presented as mean ± SD. n>3. *P < 0.05 versus BAP31^-/-^, **P < 0.01 versus BAP31^-/-^, ***P < 0.001 versus BAP31^-/-^.#P < 0.05 versus con group, ##P < 0.01 versus con group, ###P < 0.001 versus con group.

### IL-4 recovered the down regulation of C/EBP β induced by BAP31 in RAW264.7 cells

To further verified the phenomena of figure 3, we established a knockdown cell line lacking BAP31 in RAW264.7 cells. Due to the low efficiency of transfecting RAW264.7 cells by lipofectamine, we used virus transfection to construct the stable cell lines with BAP31 knockdown. The mRNA and protein expression of BAP31 in RAW264.7 cells were reduced about 70% compared with that of control cells (**Fig 4A and B**). Similar to the studies on BMDMs, we evaluated the impact of BAP31 knockdown on the expression of C/EBP β in RAW264.7 cells under various conditions. As expected, the results of RAW264.7 cells are consistent with those of BMDMs (**Fig 4C-G**).

**Figure 4.**
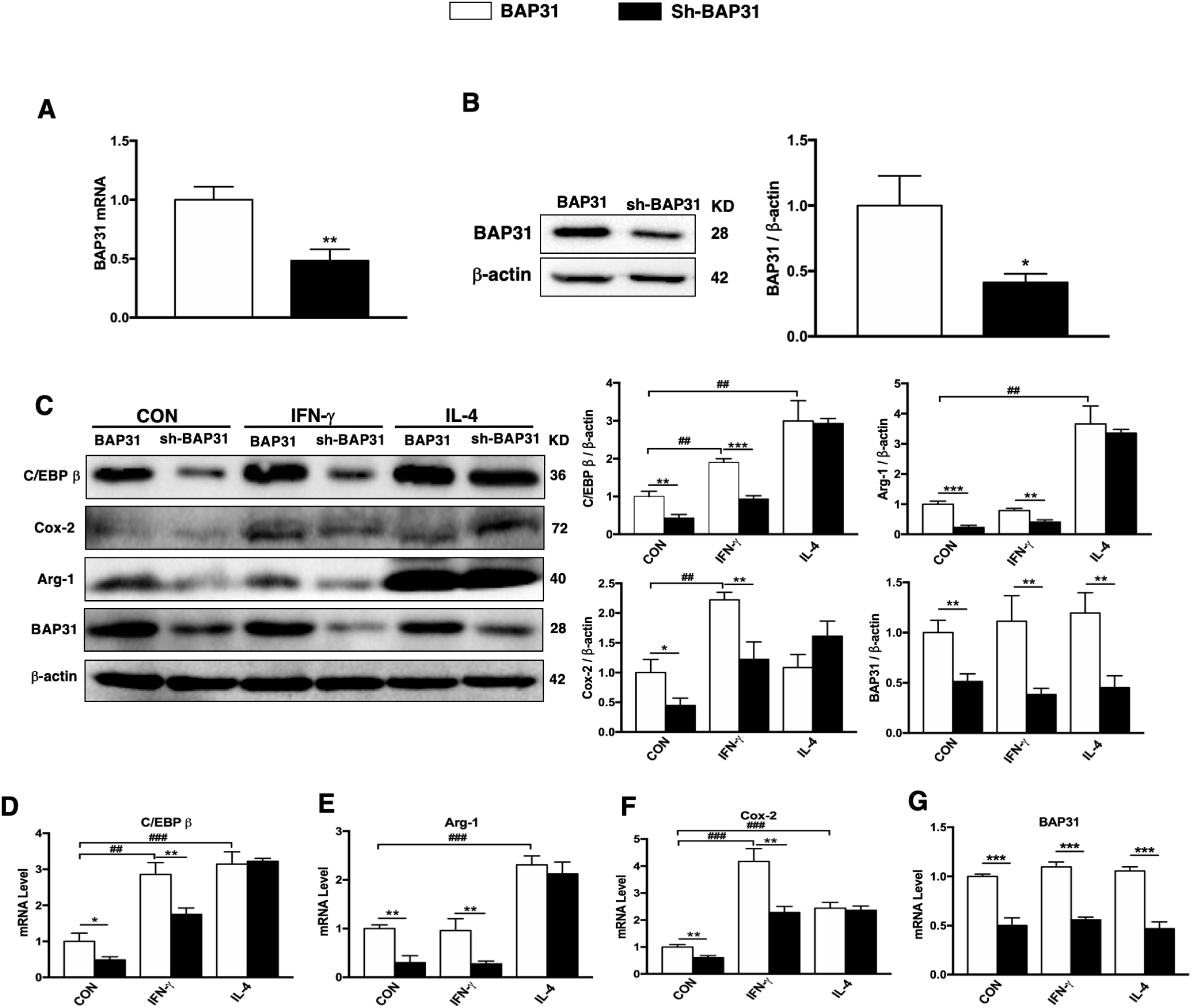
IL-4 recovered the down regulation of C/EBP β induced by BAP31 in RAW264.7 cells. A RT-qPCR analysis of BAP31 mRNA from RAW264.7 cells and Sh-BAP31-RAW264.7 cells. B Western-blot analysis of BAP31 protein expression in RAW264.7 cells and Sh-BAP31-RAW264.7 cells, histograms showed BAP31 relative change. C Western-blot analysis of C/EBP β, Cox-2 and Arg-1 protein expressions in RAW264.7 cells and Sh-BAP31-RAW264.7 cells, samples were divided into con group, IFN-γ (20 ng/mL) stimulated group and IL-4 (20 ng/mL) stimulated group, both stimulated for 24 h. The histograms showed the relative changes. D-G RT-qPCR analysis the C/EBP β, Cox-2 and Arg-1 mRNA of the above cells. Data information: In (A-G) data presented as mean ± SD. n>3. *P < 0.05 versus BAP31^-/-^, **P < 0.01 versus BAP31^-/-^, ***P < 0.001 versus BAP31^-/-^.##P < 0.01 versus con group, ###P < 0.001 versus con group.

### BAP31 regulated C/EBP β through Egr-2

To clarify the mechanisms underlying that BAP31 regulated the expression of C/EBP β and M2 macrophages, we first examined the association between the two proteins, and the results of the immunoprecipitation showed that there was no binding between BAP31 and C/EBP β (**Fig EV3A**). Therefore, BAP31 regulated C/EBP β was not mediated through a direct interaction. Then our attention was drawn to CREB and Egr-2 which could regulate the expression of C/EBP β. The results demonstrated that there was no significant change of CREB in BAP31^-/-^ group, but the Egr-2 protein was consistent with the change of C/EBP β (**Fig 5A**). Then, we detected whether the mRNA level of Egr-2 was influenced by BAP31. Interestingly the mRNA of the Egr-2 remained unchanged (**Fig EV3B**), which indicating that BAP31 deficiency decreased the protein level of Egr-2 might through posttranscriptional mechanisms. Therefore, we suspected that BAP31 modulated C/EBP β maybe through regulating the protein expression of Egr-2. To demonstrate this hypothesis, we designed the over expression plasmid of Egr-2 and transferred it into BAP31^-/-^ BMDMs. Western blot results revealed that the expression of Egr-2 was increased after transferred the plasmid (**Fig EV3C**). Subsequently, we detected the protein levels of C/EBP β and found that Egr-2 overexpression significantly inhibited the decreased of C/EBP β in BAP31^-/-^ BMDMs. The C/EBP β expression levels remained the same with the BAP31^+/+^ BMDMs group (**Fig 5B**). That is to say overexpression of Egr-2 restores the decrease of C/EBP β caused by BAP31 knockdown. These data implied that BAP31 regulated C/EBP β may be through Egr-2.

**Figure 5.**
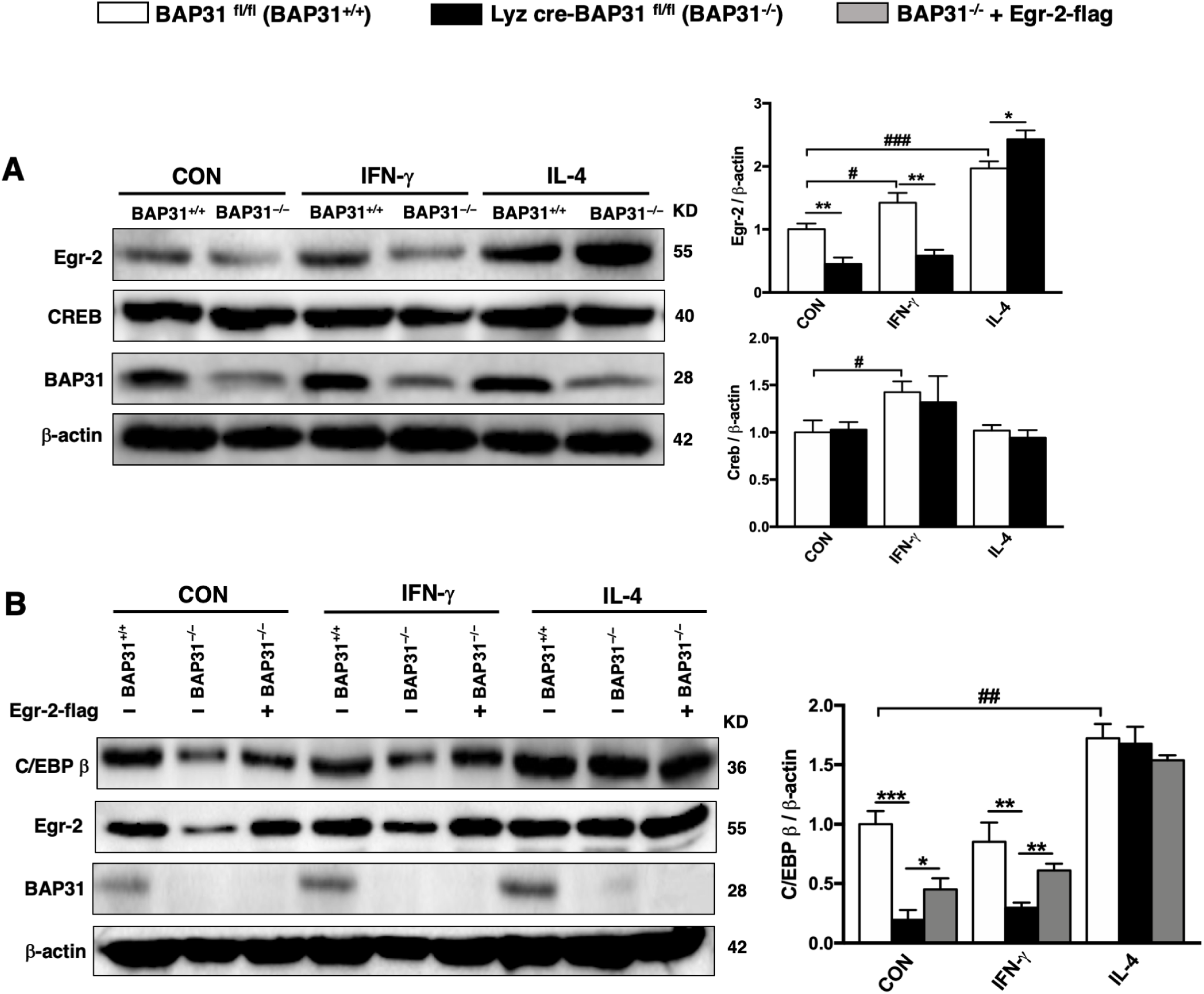
BAP31 regulated C/EBP β through Egr-2. A Western-blot analysis of Egr-2 and CREB protein expressions in BMDMs of BAP31^-/-^ and BAP31^+/+^ mice, samples were divided into con group, IFN-γ (20 ng/mL) stimulated group and IL-4 (20 ng/mL) stimulated group, both stimulated for 24 h. The histograms showed the relative changes. B Western-blot analysis of C/EBP β protein expressions in BMDMs of BAP31^+/+^, BAP31^-/-^ and BAP31^-/-^ + Egr-2-flag, samples were divided into con group, IFN-γ (20 ng/mL) stimulated group and IL-4 (20 ng/mL) stimulated group, both stimulated for 24 h. The histograms showed the relative changes. Data information: In (A, B) data presented as mean ± SD. n>3. *P < 0.05 versus BAP31^-/-^, **P < 0.01 versus BAP31^-/-^, ***P < 0.001 versus BAP31^-/-^.#P < 0.05 versus con group, ##P < 0.01 versus con group, ###P < 0.001 versus con group.

### BAP31 regulated C/EBP β through IL-4/IL-4Rα/Egr-2 signal pathways in IL-4-stimulated macrophages

BAP31 regulated C/EBP β through Egr-2, however, the connection to IL-4 signal pathway is unknown. We next sought to explore why IL-4-stimulation could recover the down regulation of C/EBP β induced by BAP31 deletion. We investigated the IL-4 specific receptor IL-4Rα in the IL-4 signaling pathway (Junttila, 2018). The results shown that the protein expression level of IL-4Rα was significantly increased in BAP31^-/-^ group (**Fig 6A**), while the mRNA expression was unchanged (**Fig EV4A**). Then we transferred the IL-4Rα siRNA into BAP31^-/-^ BMDMs, and found that IL-4Rα was decreased (**Fig EV4B**). Subsequently, after IL-4Rα knockdown, resulted in IL-4-stimulation could not recover the down regulation of C/EBP β induced by BAP31 deletion (**Fig 6B**). We came to a conclusion that the regulation of BAP31 on C/EBP β related to the expression of IL-4Rα in IL-4-stimulated macrophages. Combined with the results of Figure 5, we doubted that BAP31 regulated C/EBP β and M2 macrophages through the IL-4/IL-4Rα/Egr-2 signal pathways. To test our suspicions, the target factors of the signal pathways were knockdown or over expressed with different combination in BAP31^-/-^ BMDMs. As expected, knockdown Egr-2 partially blocked the IL-4 induced recovery of C/EBP β expression while knockdown both Egr-2 and IL-4Rα fully blocked the IL-4 induced recovery of C/EBP β expression. Interestingly, after simultaneously over expressed Egr-2 and knockdown IL-4Rα, the expression of C/EBP β could return to the original level stimulated by IL-4 (**Fig 6C and EV4C**). Those findings indicated that BAP31 regulated C/EBP β and M2 macrophages through Egr-2 and IL-4Rα in IL-4/IL-4Rα/Egr-2 signaling pathways.

**Figure 6.**
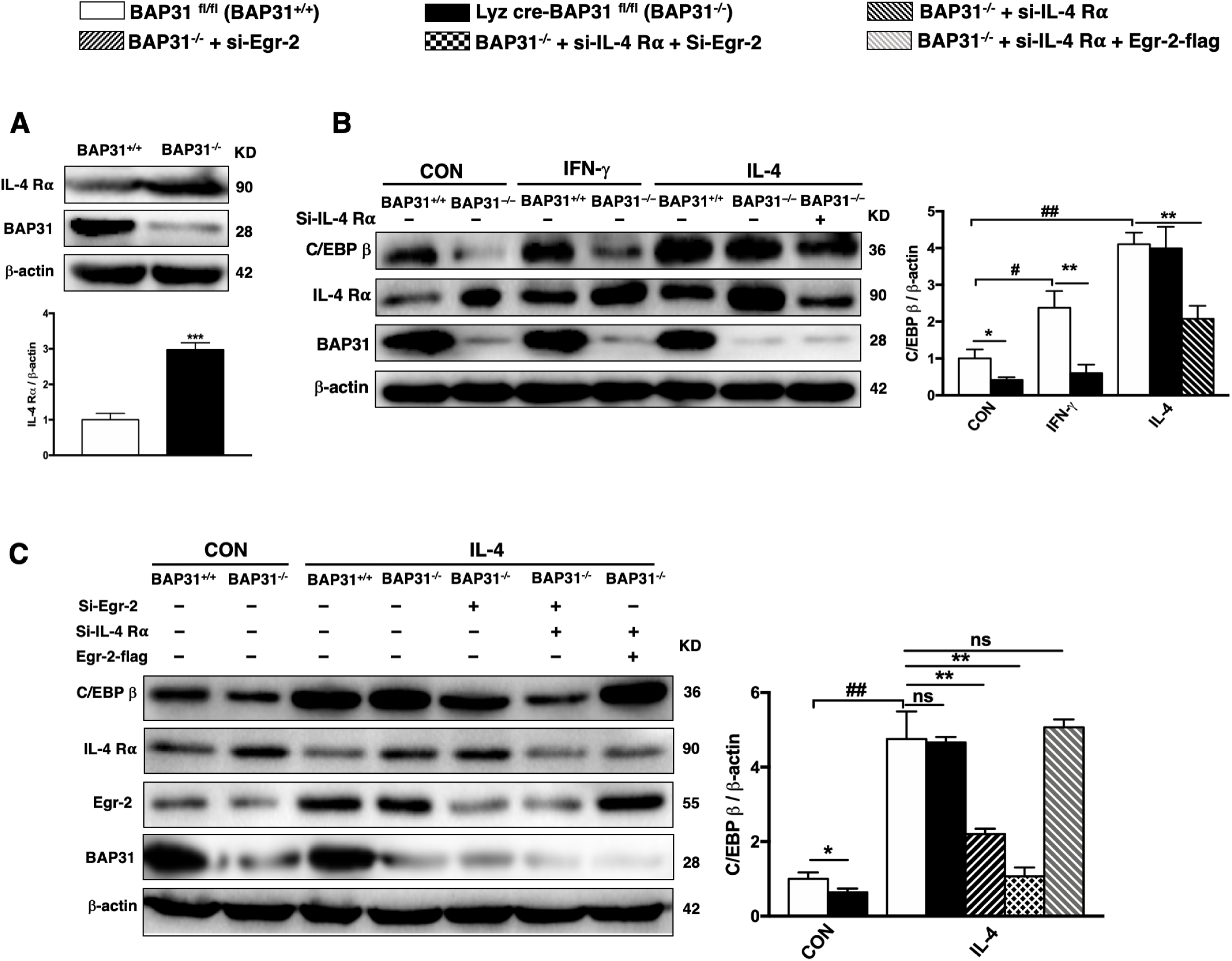
BAP31 regulated C/EBP β through IL-4/IL-4Rα/Egr-2 signal pathways in IL-4-stimulated macrophages. A Western-blot analysis of IL-4Rα protein expressions in BMDMs of BAP31^-/-^ and BAP31^+/+^ mice, histograms showed the relative changes. B Western-blot analysis of C/EBP β protein expressions in BMDMs of BAP31^+/+^, BAP31^-/-^ in con group and IFN-γ (20 ng/mL) stimulated group, and BAP31^+/+^, BAP31^-/-^, BAP31^-/-^ + si-IL-4Rα, in IL-4 (20 ng/mL) stimulated group. The histograms showed the relative changes. C Western-blot analysis of C/EBP β protein expressions in BMDMs of BAP31^+/+^, BAP31^-/-^ in con group, and BAP31^+/+^, BAP31^-/-^, BAP31^-/-^ + si-Egr-2, BAP31^-/-^ + si-IL-4Rα + si-Egr-2, and BAP31^-/-^ + si-IL-4Rα + Egr-2-flag, in IL-4 (20 ng/mL) stimulated group. The histograms showed the relative changes. Data information: In (A-C) data presented as mean ± SD. n>3. *P < 0.05 versus BAP31^-/-^, **P < 0.01 versus BAP31^-/-^, ***P < 0.001 versus BAP31^-/-^.#P < 0.05 versus con group, ##P < 0.01 versus con group.

### Depiction of BAP31 regulated polarization of macrophages through C/EBP β in cutaneous wound healing

The functions of the IL-4/IL-4Rα signaling pathway have been extensively studied in recent years. This pathway plays an indispensable role in M2 polarization of macrophages. When IL-4Rα on macrophage surface combines with IL-4, it will activate its downstream proteins, and result in macrophages secrete CD206, Arg-1 and other cytokines to play anti-inflammatory functions. Our findings suggested that BAP31 interferes the expression of IL-4Rα and Egr-2 in the IL-4/IL-4Rα/Egr-2 signaling pathways. BAP31 positively regulated Egr-2, influenced C/EBP β and its downstream cytokines, and then affected the repair of skin injury. At the same time, BAP31 negatively regulated the expression of IL-4Rα, and leaded to recover the effect of BAP31 on C/EBP β expression stimulated by IL-4 (**Fig 7**). As depicted in Figure 7, BAP31 regulated polarization of macrophages and C/EBP β through IL-4Rα and Egr-2 in cutaneous wound healing.

**Figure 7.**
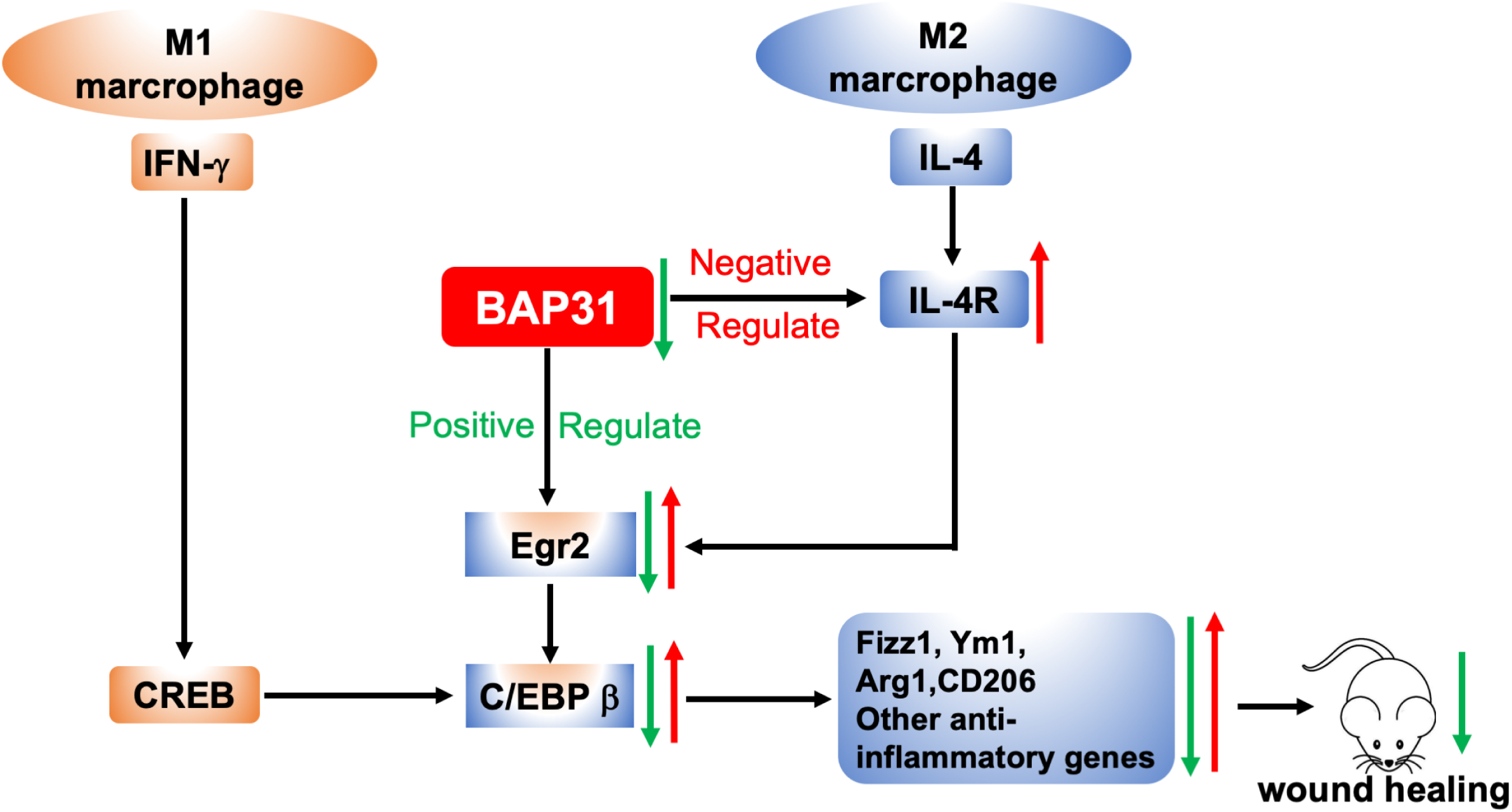
Depiction of BAP31 regulated polarization of macrophages through C/EBP β in cutaneous wound healing. Graphical summary, the pathways indicated by the green arrow represented the positive regulation of BAP31 on C/EBP β, the absence of BAP31 causes the decrease of C/EBP β through Egr-2, which in turn affected the M2 polarization of macrophages and the cutaneous wound healing. The pathways indicated by the red arrow represented the nagative regulation of BAP31 on IL-4Rα with IL-4 stimulated, the change caused by the green arrow pathway was restored due to the increased of IL-4Rα after BAP31 knockout. **Expanded View**

## Discussion

In adult wound healing, macrophages play key roles in all phases, including inflammation, proliferation, and remodeling (Munir *et al*, 2020). In wounds heal process, the local macrophage population transitions from predominantly pro-inflammatory (M1-like phenotypes) to anti-inflammatory (M2-like phenotypes) (Krzyszczyk *et al*, 2018). Nevertheless, the role of BAP31 in wound healing and M2 polarization of macrophages have not been reported.

B cell receptor associated protein 31 (BAP31) is a mainly expressed protein of the ER and part of a large BAP family that interacts with mIgD, cellubrevin, major histocompatibility complex class I (MHC I), BCL-2/BCL-X(L), or membrane-associated RING-CH (MARCH), and plays an important role in regulating the intracellular transport of these proteins (Means *et al*, 2010; Song C *et al*, 2009). Additionally, the closely related MARCH-I and MARCH-VIII were found to ubiquitinate and down-regulates MHC-II, a protein that widespread on macrophage surface to participate in immune response (Liu H *et al*, 2016). Moreover, BAP31 also affected the activation and secretion of cytokines of T cells that could influenced macrophages (Niu *et al.*, 2017). All of these findings led us to speculate that BAP31 may be closely related to the immune function of macrophages, and likely to affect the immune response of macrophages related to MHC-II and participate in the transformation of M0 macrophages to M1 and M2 macrophages. However, the role of BAP31 in macrophages has not been investigated. Therefore, we generated the mice which BAP31 conditional knockout in macrophages.

Macrophages were immune cells with various functions, including phagocytosis and cellular immunity. In immune repair, macrophages can engulf and destroy the injured tissue, which is helpful to start the recovery process(Esposito *et al.*, 2019; Kim *et al.*, 2019). In order to test our conjecture that BAP31 may affect the functions of macrophages, we used BAP31 conditional knockdown mice and normal mice to create cutaneous wound models, and observed that BAP31 knockout significantly slowed down the repair process. In the later course of healing, the expression of Ym1, an important anti-inflammatory protein, was also significantly reduced after BAP31 knockdown. It has been reported that during the healing process, abundant apoptotic debris and anti-inflammatory cytokines induced the production of M2 macrophages, which contribute to tissue regeneration and repair (Engel JE & Chade AR, 2019; Zhang *et al.*, 2018). Therefore, we speculated that the effect of BAP31 on wound healing might related to M2 polarization of macrophages.

To test this hypothesis, we used BMDMs from BAP31 conditional knockout mice and RAW264.7 cells which BAP31 knockdown achieved via shRNA to identify the markers involved in M2 macrophages. We found that BAP31 knockdown reduced the expression of M2 macrophage markers such as CD206, Ym1, Arg-1 and Fizz1 (Montoya *et al*, 2019). One of the possible explanations for how BAP31 regulated these genes was that BAP31 is an important transporter regulatory protein could affect the activities of transcription factors of M2 macrophage markers (Sun *et al*, 2017). Our further investigations confirmed this speculation and demonstrated that the deletion of BAP31 will result in the decrease of many transcription factors related to M2 macrophages contained IRFs, C/EBPs, PPAR-γ, and the most obvious was C/EBP β. In order to confirm the regulation of BAP31 on C/EBP β, we overexpressed BAP31 in RAW264.7 cells, and found thatthe expression levels of C/EBP β and M2 macrophage markers were increased. We came to the conclusion that BAP31 affected M2 macrophages through the positive regulation of C/EBP β both in BMDMs and RAW264.7 cells.

To clarify the underlying mechanism that BAP31 regulated C/EBP β, we performed the IP analysis and found that there was no direct binding between BAP31 and C/EBP β. Therefore, we considered that BAP31 may regulated some factors to affect C/EBP β production. Regulatory factors affecting C/EBP β transcription containe Sp1, CREB, SREBP1c, RARα, Myβ, Fra-2, Egr-2, STAT-3, NFκB. We focused on CREB and Egr-2, which were closely related to M2 macrophages (Chen Z *et al*, 2005; Lamkin DM *et al*, 2019; van der Krieken *et al*, 2015). There were two cAMP-like responsive elements (CRE-like sites) in the region close to the TATA box of the C/EBP β gene and the PKA/CREB pathway targets these two CRE-binding sites, and thereby regulates the transcription of C/EBP β (Niehof *et al*, 2001). However, the detailed ways of Egr-2 regulating C/EBP β have not been reported, maybe related to “leaky ribosome scanning mechanism” (Kozak M, 1989). Our results showed that the expression of Egr-2 protein decreased significantly after BAP31 knockdown and when we restored the level of Egr-2 resulted in the recovery of the down regulation of C/EBP β induced by BAP31 deficiency. Therefore, we concluded that the regulation of BAP31 on C/EBP β was through Egr-2, a transcript factor can affect the expression of C/EBP β. Because BAP31 knockdown did not affect the expression of Egr-2 mRNA, we believed that BAP31 only affected the degradation of Egr-2 protein. Early growth response 2 (Egr-2) was part of transcription factors Egr family, could be degradated by ubiquitin-dependent proteasome system. BAP31 regulated the degradation of Egr-2 possibly through ubiquitin E3 ligase which were participated in the degradation of Egr-2 such as WWP2, AIP2, CbI-b (Chen *et al*, 2009; Chen *et al*, 2014; Safford *et al*, 2005). However, the detailed mechanisms and contributions remain to be clarified.

Macrophages could be polarizated by IFN-γ and IL-4 to show classical activation (M1) and alternative activation (M2) respectively, and C/EBP β can also increase after macrophages activated, so we tested the effect of BAP31 on C/EBP β expression in macrophages after treated with IFN-γ and IL-4 (Goodman S *et al*, 2019; Veremeyko *et al.*, 2018). Our results suggested that the effect of BAP31 on C/EBP β was restored in macrophages treated with IL-4. Further studies revealed that not only C/EBP β, and its downstream transcript factors Cox-2 and Arg-1 were recovered after IL-4-stimulated. We speculated that interfered factors in the IL-4 signal pathways might counteract the decrease of C/EBP β induced by BAP31 knockdown. Previous studies had demonstrated that when IL-4 upon binding to receptors (IL-4Rα1, IL-13Rα1, or IL-13Rα2), JAK1 and JAK3 are activated, leading to activation of STAT6 and IRF4, which is the classical signal pathways of IL-4 (Essandoh K *et al*, 2016). Based on this, we first detected the upstream protein IL-4Rα, and the results revealed that BAP31 knockdown up regulated the expression of IL-4Rα. Interestingly knocked down IL-4Rα blocked the IL-4-induced the recovery of the C/EBP β expression. Therefore, we inferred that after BAP31 knockdown, the increased expression of IL-4Rα affected its downstream factor C/EBP β. Our results had proved that BAP31 affected the expression of C/EBP β through Egr-2, then regulated the M2 polarization of macrophages and cutaneous wound healing. Furthermore, BAP31 can regulated the expression of IL-4Rα, and then affected the changes of Egr-2 and C/EBP β through IL-4/IL-4Rα/Egr-2 signaling pathways under IL-4-stimulation.

In summary, these findings demonstrated that BAP31 regulated C/EBP β through Egr-2 and IL-4Rα and then affected M2 polarization of macrophages and cutaneous wound healing. The data presented here illustrate a new potential function of BAP31 and provide novel targets for the prevention and treatment of chronic wounds.

## Materials and methods

### BAP31 conditional knock out mice

BAP31^flox/flox^ mice were reported previously by our laboratory. BAP31^flox/flox^ mice were mated with age-matched Lyz2-cre transgenic mice to generate Lyz2-cre BAP31^flox/flox^ and BAP31^flox/flox^ mice. Gender same and littermates Lyz2-cre BAP31^flox/flox^ and BAP31 ^flox/flox^ mice were used throughout all experiments. All mice were housed under specific pathogen free conditions at room temperature under a 12 h light/dark cycle, and fed with mouse nutritional food. Animal experiments were approved by the Institutional Animal Care and Use Committee of Northeastern University, and all mice were treated in accordance with the Guide for the Care and Use of Laboratory Animals.

### Cell culture and reagents

Mouse bone marrow derived macrophages (BMDMs) were extracted from mice of 6-9 wk of age, cultured in Roswell Park Memorial Institute (RPMI) Medium (Gibco BRL Life Technologies, CA, USA) with10% (v/v) fetal bovine serum (FBS), penicillin (100 U/mL) and streptomycin (100 μg/mL) and 10% L929 cellular supernatants (Racioppi *et al*, 2019). BMDMs were maintained 7 days for future experiments. RAW 264.7 mouse mononuclear macrophage leukemia cell line was obtained from the American Type Culture Collection (Manassas, VA, USA). The cells were maintained in Dulbecco’s modified Eagle Medium (DMEM) (Gibco BRL Life Technologies, CA, USA) with GlutaMAX TM containing 10% (v/v) fetal bovine serum (FBS), penicillin (100 U/mL) and streptomycin (100 μg/mL). The cultures were maintained at 37 °C in a humid incubator with a 5% (v/v) CO_2_ atmosphere.

### Virus packaging and cell line transfection

For the transfection, 3×10^6^ cells were seeded in 60 mm dish 24 h prior to transfection in DMEM supplemented with 10% FBS and 1% penicillin/streptomycin. For knockdown, the cells were transfected with total of 5 μg of plasmid DNA (1.5 μg PSPAX2, 1.5 μg PMD_2_G, and 2 μg PL/shRNA/GFP-mouse-Bap31) in a volume of 100 μL DMEM medium without FBS was mixed with 10 μL Lipofectamine (Invitrogen, CA, USA). For overexpression, the cells were transfected with total of 10 μg of plasmid DNA (2.5 μg PLP1, 1.75 μg PLP2, 2.5 μg PCMV and 3.25 μg pLBap31-flag or 3.25 μg pLEgr-2-flag) in a volume of 100 μL DMEM medium without FBS was mixed with 10 μL Lipofectamine. Cells transfected with plasmid vector were used as control. 48 h and 72 h after transfection, we collect cell supernatant separately for next use. 293T cells supernatant with the packaged virus was collected and filtered through 0.45 μM cellulose acetate filters, then transduce 1×10^6^ RAW264.7 Cells or BMDM cells in a 6-well plate. 72 h post-transduction, cells were observed using Leica DMI3000 B fluore scence microscope (Wetzlar, Germany). Then, 10 μg/mL blasticidin and the limited dilution method were used to obtain the monoclonal cell strains, and the transduction efficiency was quantified using Western blot.

### Small interfering RNA (siRNA) transfection

A total of 1×10^6^ BMDM cells were seeded in a 6-well plate, then transfected siRNA (50 nm) mixed with Lipofectamine 3000 according to the instructions. The negative siRNA was used to generate the control cells. After transfection for 72 h, Real-time PCR and Western blot were used to detect the transcription levels and protein expression of the targets. The sequences of the siRNA used in this study were as follows: siRNA-Egr-2 sense, 5’-CCUCGAAAGUACCCUAACATT-3’ and siRNA-Egr-2 antisense 5’-UGUUAGGGUACUUUCGAGGTT-3’; siRNA-IL-4Rα sense, 5’-GCCAGGAGUCAACCAAGUATT-3’ and siRNA-IL-4Rα antisense, 5’-UACUUGGUUGACUCCUGGCTT-3’ (GenePharma, Shanghai, China).

### Fluorescence-activated cell sorter (FACS) analysis of BMDMs

BMDMs were collected and cell concentration was adjusted to 5×10^4^/100 µL. After cells were washed twice with cold PBS with 2% BSA, Fc receptor block (Miltenyi Biotec Inc) was added and incubated on ice-water bath for 10 min. Wash once with cold PBS with 2% BSA, followed by FITC-CD11b (BD, NY, USA), and PE-F4/80 (BD, NY, USA) mAb were added to the cells which were then incubated on ice water bath at 4 °C for 45 min, and washed with PBS with 2% BSA. Then analyzed using Fortessa or Accuri C6 and software (BD, NY, USA).

### Reverse transcription and quantitative real-time PCR (RT-qPCR)

Total cellular RNA was isolated from the above two cells using TRIzol reagent (Thermo Fisher Scientific, MA, USA) according to the manufacturer’s instructions. The RNA samples (2 μg) were used for the synthesis of cDNA by reverse transcription using GoScript ™ Reverse Transcriptase (Promega, Madison, USA). RT-PCR was performed with a CFX96 Touch ™ Real-Time PCR Detection System (Bio-Rad Laboratories). The PCR mixture was at a volume of 10 μL containing 5 μL SYBR Premix Ex Taq (Promega, Madison, USA), 0.5 μM each of the primers, and 1 μL cDNA prepared as described and using the following conditions: 95 °C for 2 min, 95 °C for 15 sec, and 60 °C for 60 sec for 40 cycles. The results were analyzed according to the 2^−ΔΔCq^ formula. The sequences of primers are listed in **Table 1.**

### Skin wound healing model

Lyz2-creBAP31^flox/flox^ and BAP31^flox/flox^ mice were injected intraperitoneally with chloral hydrate (4%, 350 mg/kg) for general anesthesia. Then a 1 cm square wound was cut on the back of the mouse with sterile eye scissors, and a 0.5 mm thick silicone film was sewn on the wound to prevent the wound from contracting. At different time points (0 days, 1 day, 4 days, 7 days, 10 days and 14 days), the wounds of mice were photographed, and the wound area was calculated by ImageJ software (NIH, Bethesda, MD). The percentage of wound area was calculated as follows: wound area (%)= (wound area on day n/wound area on day 0) × 100%.

### Histology analysis

The mice were euthanized at day 14 after injury, and skin tissue samples were harvested and fixed with Paraformaldehyde (4%), embedded in paraffin, and sequentially sectioned. Hematoxylin-Eosin (H&E) was performed to detect skin form and wound healing. Immunofluorescence staining was carried out to determine the anti-mouse Ym1 (STEM CELL, CA, USA) at the injured tissue paraffin slides, and the tissue samples was also stained with 4’, 6-diamidino-2-phenylindole (DAPI).

### Cytokines measurement using FACS

The collected cells were suspended in PBS, fixed in 1 mL 2% paraformaldehyde on ice for 1 h. Fc receptor blocker (Miltenyi Biotec Inc) was added and ice-water bath was used for 10 minutes. Wash once with cold PBS, followed by FITC-CD11b (BD, NY, USA), and PE-F4/80 (BD, NY, USA) mAb were added to the cells, then cells were suspended in 100 μL of 0.1% saponin (Sigma, MO, USA) mixed with 1:1000 each of purified anti-Ym1 rabbit antibody, anti-CD206 rabbit antibody (EB, MA, USA) or anti-C/EBP β rabbit antibody (abcam, MA, USA). After washing, samples were subjected to analysis on an FACS Accuri C6 (BD, NY, USA) and analyzed using FlowJo 7.6 software (TreeStar, Ashland, OR).

### Western blot analysis

Proteins were isolated from the cells in buffer RAPI (50 mM Tris-Hcl, pH 7.4, 0.2% Triton-X-100, 0.1% SDS, 150 mM Nacl, 2 mM EDTA, 50 mM NaF, 10% Na-deoxycholate, 2.5 mM sodium pyrophosphate decahydrate, 1 mM sodium orthovanaldate, 1 mM PMSF) at 4 °C for 30 min, centrifuged at 12000 g/15 min, collect supernatant add 5×sodium dodecyl sulfate (SDS) sample buffer, and boiled 10 min. Lysates (20 μg) were electroblotted onto a polyvinylidene difluoride membrane (Milipore, MA, USA) following electrophoretic separation in 10% or 12% SDS-polyacrylamide gel, Bloked with 5% BSA-TBST (Tris-buffered saline/Tween 20) at room temperature for 1 h, and followed by incubation overnight at 4 °C with the primary antibodies, And then washed with TBST three times, incubated with the secondary antibody for 1 h at room temperature The signal was detected with ECL (Tanon, Shanghai, China). Protein patterns were documented on a chemiDoc XRS+ System (Bio Rad, Munich, Germany). and Optical density analysis of signals was performed with Image Lab software (Bio Rad, Munich, Germany). The antibodies source was listed: anti-BAP31, anti-C/EBP β, anti-CREB, anti-Egr-2, anti-IL-4Rα were from abcam (Cambridge, MA, USA); anti-Arg-1, anti-Cox-2 were from Cell Signaling Technology (Danvers, MA, USA); anti-Ym1 was from STEM CELL (STEM CELL, CA, USA). The band intensities of the target proteins were calculated and normalized to that of β-actin.

### Coimmunoprecipitation

Cells were treated with RIPA lysis buffer (Beyotime Biotechnology, Shanghai, China) and mixed with protein G (Beyotime Biotechnology, Shanghai, China) for 30 min. The lysates were incubated with antibodies at 4 °C for 2 h, then washed the complexes bound to protein G three times and eluted in 5×SDS sample buffer. All samples were continue analyzed by Western blot analysis.

### Statistical analysis

All experiments were performed at least three to five times, and all experimental data are reported as means ± experimental standard deviations (SD). Student’s t-test was used to determine the significance of the results (significance: *P < 0.05; **P < 0.01; ***P < 0.001).

## Abbreviations

BAP31: B cell receptor associated protein 31;
ER: endoplasmic reticulum;
BMDM: bone marrow derived macrophage;
Arg-1: arginase-1;
CD206: cluster of differentiation 206;
Ym1: Chitinase-like 3;
Fizz1: Resistin like alpha;
DAPI: 4’, 6-diamidino-2-phenylindole;
HE: hematoxylin and eosin;
IFN-γ: interferon-γ;
IL-4: interleukin-4;
IL-4Rα: interleukin-4 Receptor alpha;
TCR: T Cells Receptor;
PPARγ: peroxisome proliferator-activated receptor gamma;
IRF: Interferon regulatory factors;
C/EBP: CCAAT-enhancer-binding proteins;
Egr-2: early growth response 2.

## Acknowledgments

This research was supported by the Liao Ning Revitalization Talents Program (XLYC 1902063) and National Natural Science Foundation of China (31370784, 31670770, 2016YFC1302402).

## Authors’ contributions

B.W. and Q.Y. designed the experiments. Q.Y. performed the experiments & data analysis, and drafted the manuscript. B.W. and XY.W. revised the manuscript. B.Z. and LJ.S. transfection over-expressed plasmid. X.L was involved in data analysis. YH.C. and Y.X discovered the change of CEBP β. The authors read and approved the final manuscript.

